# Extending scope and power of circular RNA research with circtools 2.0

**DOI:** 10.1101/2025.02.16.638209

**Authors:** Shubhada R. Kulkarni, Christoph Dieterich, Tobias Jakobi

## Abstract

Circular RNAs (circRNAs) play a pivotal role in gene regulation, acting as transcriptional modulators at various cellular levels and exhibit remarkable specificity across cell types and developmental stages. Detecting circRNAs from high-throughput RNA-seq data presents significant challenges, as it requires specialized tools to identify back-splice junctions (BSJ) in raw sequencing data. To address this need, we previously developed *circtools*, a robust and user-friendly software for circRNA detection, downstream analysis, and wet-lab integration. Building on this foundation, we now introduce *circtools 2.0*, a significantly enhanced framework designed to advance computational and support experimental analysis for circular RNA research. Novel key features of *circtools 2.0* include: (1) an ensemble approach to circRNA detection with improved recall rates, (2) a module for automated design of circRNA padlock probes for the Xenium spatial trancriptomics platform, (3) circRNA conservation analysis across animal model species, and (4) support for the Oxford Nanopore sequencing platform for full-length circRNA detection.

**Availability:** circtools 2.0 is available at github.com/jakobilab/circtools and licensed under GPLv3.0.

## 1. Introduction

Circular RNAs (circRNAs) are a unique class of RNA molecules, which originate from linear transcripts through back-splicing Sanger *et al*. (1976). In this process, a downstream splice donor site is joined to an upstream splice acceptor site, forming a covalently closed circular structure. CircRNAs are ubiquitously expressed across a broad spectrum of species, ranging from viruses to mammals, and their prevalence underscores their evolutionary conservation and potential functional importance.

*Circtools* is as a comprehensive software pipeline that offers workflows for the detection, quality control, and downstream analysis of circRNA (Jakobi *et al*., 2019) and found widespread use in scientific projects and training courses. Based on community feedback, we focused on the development of four different features of computational circRNA analysis identified as critical for the development of the field. Specifically, we implemented new modules that address the need for an accurate localization of circRNAs in cellular or tissue contexts using specific probes for back-splicing junctions, integrated analysis of circRNA conservation between model species, combining results of selected circRNA detection algorithms for improved recall rates, and allowing for detection of full-length circRNAs from Oxford Nanopore third-generation sequencing experiments.

By integrating these novel features, we are confident that *circtools 2.0* will provide researchers with a state-of-the-art platform to explore the multifaceted roles of circRNAs in diverse biological systems (Fig. 1A).

**Fig. 1:**
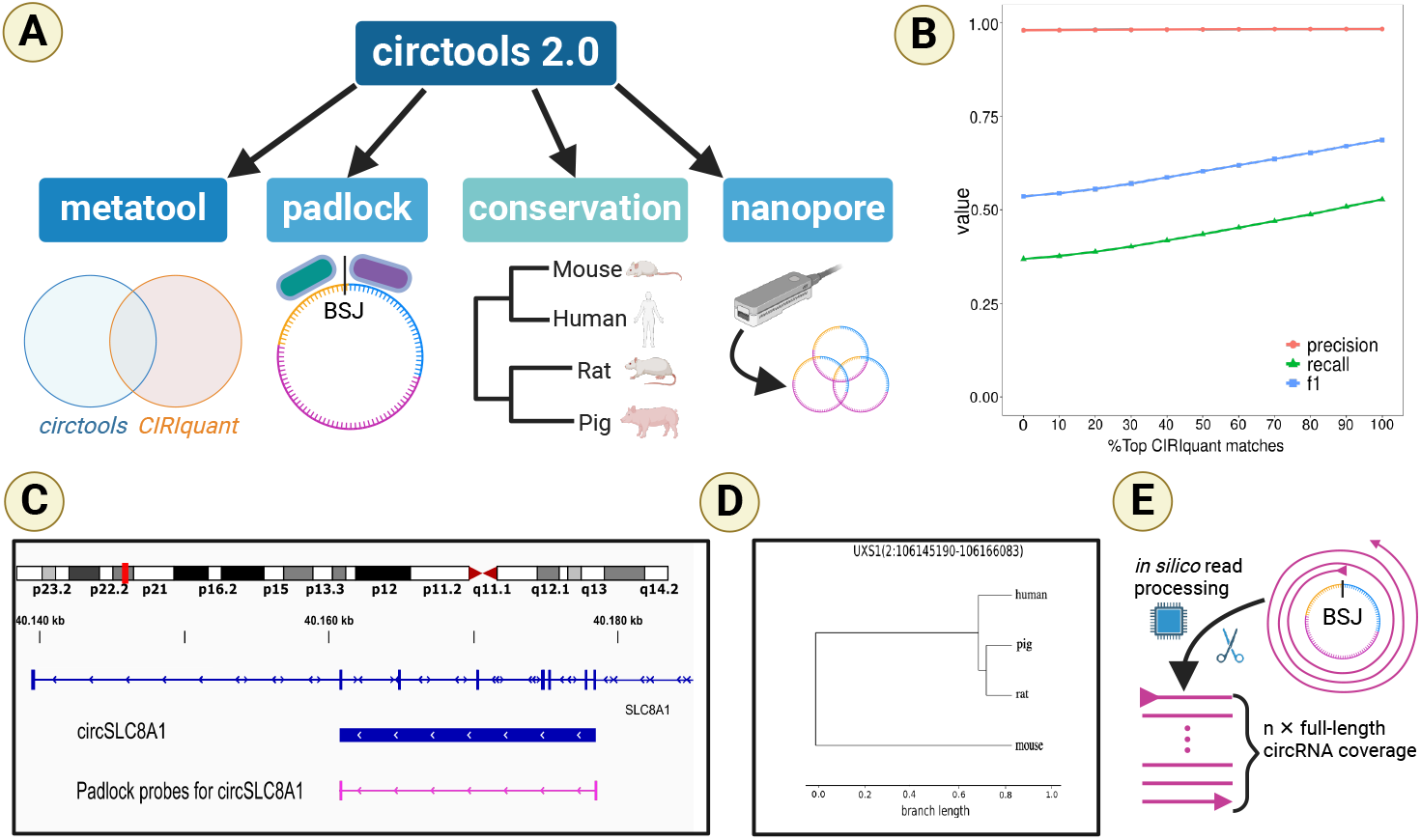
Circtools 2.0 and its new modules. **(A)** Overview of new modules introduced with *circtools 2.0*. **(B)** Precision-recall curve for *circtools detect* and *CIRIquant*. Performance measures were evaluated by adding top-scoring 10% matches from *CIRIquant* to all matches from *circtools* in increments. **(C)** Example of spatial padlock probes designed for circSLC8A1. **(D)** Example conservation plot for circUXS1 showing relative similarity in 4 species. **(E)** Simplified worklfow of the nanopore module for full-length circRNAs processing long reads, spanning the circRNA multiple times. Created with BioRender.com.

## 2. Methods

### 2.1. Metatool module

We evaluated the performance of circRNA detection tools by comparing results from the previous *circtools* version and two other popular circRNA prediction algorithms, *CIRIquant* (Zhang *et al*., 2020) and *CirComPara2* (Gaffo *et al*., 2022). The landmark study by Vromman *et al*. (2023) demonstrated that these three methods exhibit different sensitivities. We selected *CIRIquant* and *CirComPara2* as they outperformed circtools alone in precision. We used data generated as part of the Vromman *et al*. (2023) study for all subsequent evaluations. Our first pairwise comparison included *circtools* and *CirComPara2*. While *circtools* predicted 15,296 unique circRNAs, *CirComPara2* predicted 10,585 (Supplementary Fig. 1A). When we combined predictions from *CirComPara2* and *circtools*, the recall improved from 36.2% to 41.4%. However, the precision decreased from 89.2% to 68.9% (Supplementary Fig. 1B), indicating that too many false positive matches were added. Furthermore, *CirComPara2* exhibited longer running times as compared to *circtools*, which led us to exclude it from further analysis. Our next pairwise comparison comprised *CIRIquant* and *circtools*. Our results showed that both, *circtools* and *CIRIquant* detected unique circRNAs, with 9.5% (n=4,504 matches) and 53.28% (n=25,182 matches), respectively. The precision of unique circRNAs detected by *circtools* was greater than 99%. We then added the top-scoring 10% matches from *CIRIquant* to *circtools*’ predictions in increments, observing that the precision remained consistently high (*>*99%) (Fig. 1B). Moreover, we noted an improvement in recall from 36.88% to 52.75%, suggesting that the false positive rate was effectively controlled by incorporating unique matches of *CIRIquant* into *circtools*’ results.

### 2.2. Padlock probe module

Spatial transcriptomics emerged as a powerful technique to map the localization of single molecules to the level of individual cells and even offer subcellular resolution. Although most of the high-throughput methods were designed with linear polyadenylated RNAs in mind, some methods could target circRNAs as well. Padlock probes are synthetic DNA sequences (single-stranded) that are designed to target specific region of interest. Due to their high sensitivity and specificity, padlock probes are getting used in various next-generation technologies. We added the *padlock* module to *circtools 2.0* to allow spatial analysis of several circRNA and corresponding linear transcripts by *in situ* hybridization in parallel. This module was specifically tailored to the Xenium platform as it offers subcellular resolution and an option for custom panel design. The module requires three inputs: 1) circRNA coordinates detected using *circtools*’ detect step, 2) a genome FASTA file, and 3) a transcriptome GTF file. Briefly, the module identifies 50-bp long target sequences that overlap with back-splice splice junctions. Subsequently, it employs a 40-bp scanning window approach to position two adjacent 20-bp padlock probes for each target sequence (Fig. 1C; Supplementary Fig. 2). We tested all designed probes as part of a custom-built gene panel comprising linear and circular RNAs on human heart samples. We evaluated our padlock probe designs by comparing them with company-designed/tested panels using the 10X Xenium platform. Intriguingly, both types of panels demonstrated similar performance on this next-generation sequencing technology platform, indicating that the probes designed using our padlock module can efficiently be used for in situ hybridization (Supplementary Fig. 3). The padlock module outputs a BED12 file containing probe coordinates, which can be visualized in a genome browser to provide a better understanding of the targeted transcripts. Additionally, the module generates a detailed HTML report that guides users in selecting suitable probe designs based on criteria such as melting temperature, GC content, and specificity (Supplementary Fig. 4). This report enables users to make informed decisions regarding probe design and optimizes the detection process.

### 2.3. Conservation module

Evolutionary conservation analysis oftentimes uncovers the potential functional relevance of circRNAs by comparing their sequence and genomic position across different organisms. We developed a *conservation* module within *circtools 2.0*, which enables users to perform circRNA conservation analysis in five widely studied animal model species: mouse, human, rat, pig, and dog. The framework of the *conservation* module was developed with the flexibility to incorporate more species in the analysis by simply adding the species to the input config file. The input for this module is a list of BSJs for circRNAs in a given species, which is then compared with either a single target species or multiple target species. The two exons closest to the respective BSJ are lifted-over to the target species to identify the ortholog of the circRNA in each species. Next, *circtools 2.0* performs either pairwise or multiple sequence alignment using MAFFT (function *MafftCommandline* in *BioPython* (Cock *et al*., 2009)) to calculate the conservation score for each target species. Conservation score (ranging between 0 to 1) is defined by distance matrix i.e. higher the distance between two species, smaller the conservation between them. This approach enables users to quantify the degree of conservation across different organisms. The *conservation* module generates a phylogenetic tree to visually represent the conservation of circRNAs across the selected species (Fig. 1D). We provide alignment results in the *CLUSTAL* as well as in the .*tree* format. In addition, we provide all lifted-over coordinates of the circRNA in the target species, allowing users to further analyze and visualize the conservation patterns. We consider this *conservation* analysis module in *circtools 2.0* as a highly relevant addition for prioritizing circRNAs for further analysis based on the conservation of their exon sequences.

### 2.4. Nanopore module

Recent advances in long-read sequencing technologies have enabled the generation of full-length circRNA sequences (Rahimi *et al*., 2021; Zhang *et al*., 2021). However, the development of computational tools to perform these analyses on Oxford Nanopore datasets is lagging behind. We improve on this status quo by adding a new workflow to *circtools 2.0* that enables users to detect circRNAs from long-read sequencing Nanopore datasets: the (*nanopore* module). This module is designed to specifically process the unique characteristics of Oxford Nanopore data, i.e. the handling of sequencing reads *>* 5kb (Fig. 1E), and provides accurate and efficient detection of circRNAs. The module is based on code developed for the circNick-LRS and circPanel- LRS sequencing protocols as developed by (Rahimi *et al*., 2021). We have thoroughly updated the original code and now support newer genome assemblies for mouse (mm10) an human (hg38) as well as additional mammalian genomes for pig (susScr11), rat (rn6, rn7), and dog (canFam6). Additional genomes can easily be added by adapting YAML-based configuration files as long as the species are supported by ENSEMBL and UCSC’s Genome Browser. The configuration files are subsequently used by the new reference data retrieval system that automatically downloads and postprocesses reference data from reliable public databases such as UCSC’s Genome Browser and ENSEMBL without the need for short-term links to private online drives. Moreover, Perl scripts were replaced by new implementations in Python3 to seamlessly integrate in the *circtools 2.0* software framework and reduce the overall software maintenance burden.

### 2.5. Native docker images

In addition to the new modules, we further simplified the installation process for *circtools 2.0* through user-friendly docker containers. We support standard amd64-based architectures, as well as arm64-based systems such as Apple silicon (i.e., M1-M4) and system on a chip (SoC) solutions, such as Rasperry Pi. Our container solution allows users to easily install and run *circtools 2.0* without the need to handle complex software dependencies such as installation of R packages or Python libraries. This renders the new *circtools 2.0* essentially operating system agnostic.

## 3. Conclusion

In summary, the *circtools 2.0* software framework introduces several enhancements that significantly expand its capabilities and usability. The *metatool* module integrates detected circRNAs from two sources: *CIRIquant* and the existing *detect* module, which yields better recall rates while keeping the same precision. The *padlock* module designs probe sets for the Xenium platform to detect circular RNAs in single cells or tissues. The *conservation* evaluates the potential functional relevance of circRNAs across different organisms by quantifying their evolutionary conservation. Finally, the *nanopore* module comprises an updated workflow for detecting full-length circRNA sequences from Oxford Nanopore datasets, addressing a critical gap in current bioinformatic tools. In general, the use of native docker images simplifies the installation process and renders *circtools 2.0* operating system agnostic. These advancements collectively empower researchers to tackle complex biological questions in circular RNA research more effectively. We are certain that *circtools 2.0* will help researchers to advance our understanding of circular RNA biology.

## 4. Author contributions statement

S.R.K and T.J. developed and tested the software. S.R.K, T.J. and C.D. read and approved the manuscript.

## 5. Data availability

Source code and documentation is available under GPL v3.0 license at https://github.com/jakobilab/circtools.

