## Supplemental Figures for "Extending scope and power of circular RNA research with circtools 2.0"

January 17, 2025

### Figure S1

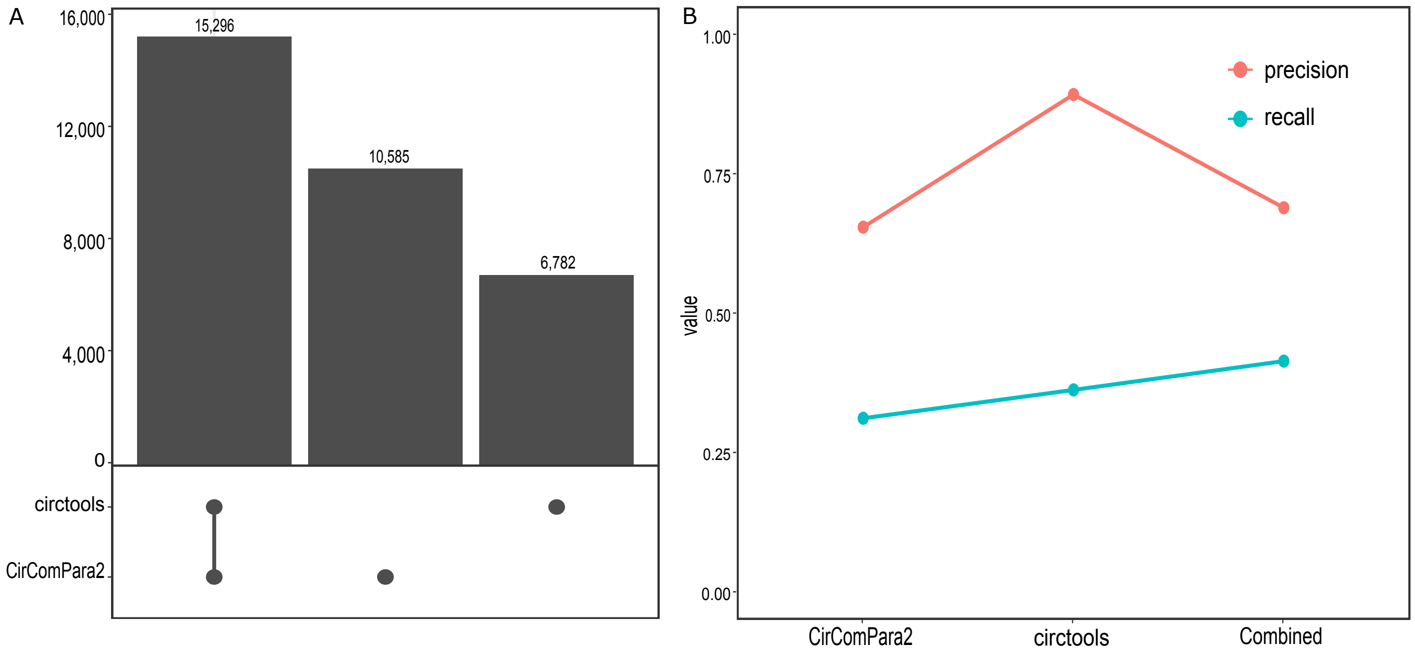


Figure 1: Common and unique matches between circtools and CirComPara2.

#### Figure S2

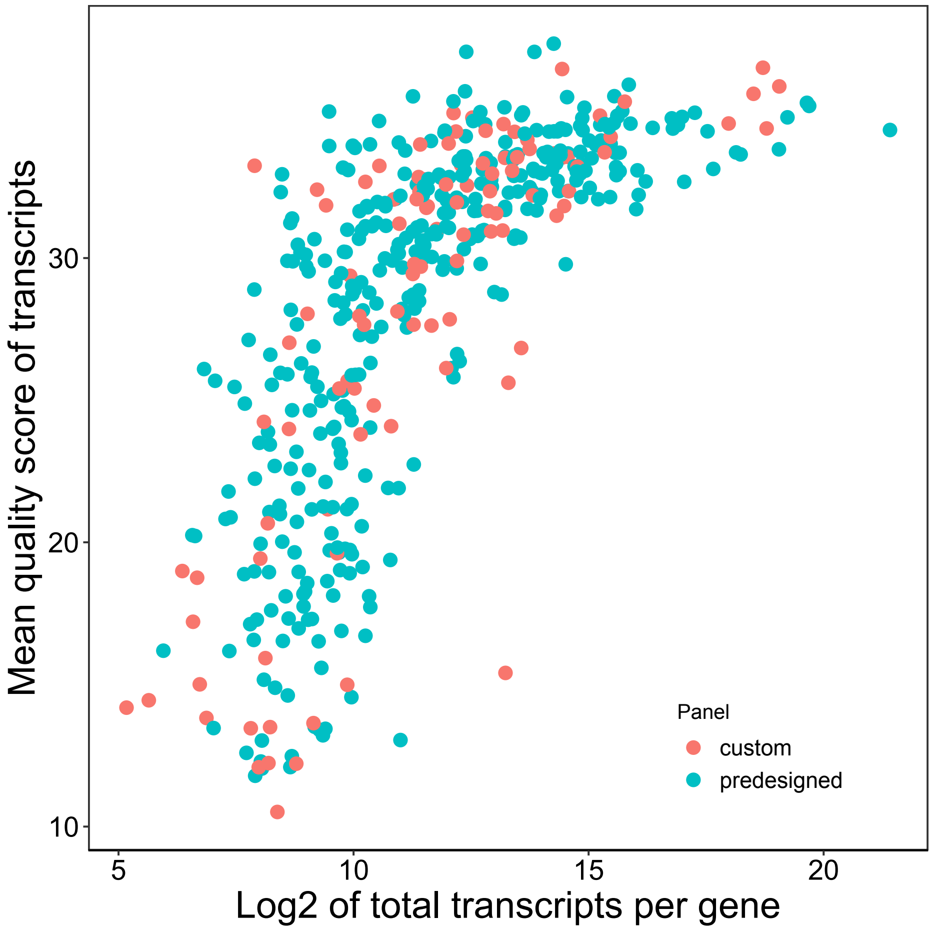


Figure 2: Distribution of quality scores for custom and predesigned probes.

### Figure S3

Colour legends for TM and GC columns

Low Medium High

| Input IDs | | | | | Designed Probes | | | | | | | | | |
|---|---|---|---|---|---|---|---|---|---|---|---|---|---|---|
| Annotation | Chr | Start | Stop | Strand | TM RBD5 | TM RBD3 | TM Full | GC% RBD5 | GC% RBD3 | Ligation Junction | RBD5 | BLAST | RBD3 | BLAST |
| UXS1 | 2 | 106093302 | 106094157 | - | 58 | 67 | 78 | 55 | 75 | preferred | TTAAACCTTCCTCCAGCGGG | 0 | ACCACGGGCTCCCACCCCAG | 0 |
| UXS1 | 2 | 106093302 | 106094157 | - | 58 | 68 | 79 | 55 | 80 | neutral | TAAACCTTCCTCCAGCGGGA | 0 | CCACGGGCTCCCACCCCAGC | 0 |
| UXS1 | 2 | 106093302 | 106094157 | - | 60 | 67 | 81 | 60 | 75 | neutral | AAACCTTCCTCCAGCGGGAC | 0 | CACGGGCTCCCACCCCAGCA | 0 |
| UXS1 | 2 | 106093302 | 106094157 | - | 62 | 66 | 81 | 65 | 70 | neutral | AACCTTCCTCCAGCGGGACC | 0 | ACGGGCTCCCACCCCAGCAT | 0 |
| UXS1 | 2 | 106093302 | 106094157 | - | 63 | 65 | 81 | 65 | 75 | neutral | ACCTTCCTCCAGCGGGACCA | 0 | CGGGCTCCCACCCCAGCATC | 0 |
| UXS1 | 2 | 106096717 | 106097710 | - | 57 | 53 | 70 | 50 | 35 | neutral | ACATCTGATGATGCTGCTGC | 0 | TGCGCAACAAGGTTTTTAAT | 0 |
| UXS1 | 2 | 106096717 | 106097710 | - | 56 | 52 | 70 | 50 | 35 | neutral | CATCTGATGATGCTGCTGCT | 0 | GCGCAACAAGGTTTTTAATT | 0 |
| UXS1 | 2 | 106096717 | 106097710 | - | 62 | 44 | 69 | 60 | 20 | neutral | TGATGATGCTGCTGCTGCGC | 0 | AACAAGGTTTTTAATTAACT | 0 |
| UXS1 | 2 | 106096717 | 106097710 | - | 62 | 45 | 69 | 60 | 25 | neutral | GATGATGCTGCTGCTGCGCA | 0 | ACAAGGTTTTTAATTAACTG | 0 |
| UXS1 | 2 | 106096717 | 106097710 | - | 62 | 45 | 69 | 55 | 25 | neutral | ATGATGCTGCTGCTGCGCAA | 0 | CAAGGTTTTTAATTAACTGA | 0 |
| UXS1 | 2 | 106096717 | 106097710 | - | 63 | 44 | 70 | 60 | 25 | neutral | TGATGCTGCTGCTGCGCAAC | 0 | AAGGTTTTTAATTAACTGAG | 0 |
| UXS1 | 2 | 106096717 | 106097710 | - | 63 | 47 | 70 | 60 | 30 | neutral | GATGCTGCTGCTGCGCAACA | 0 | AGGTTTTTAATTAACTGAGC | 0 |

Figure 3: Screenshot for padlock probe module output.
